# Lack of *Period1* accelerates colorectal tumorigenesis in Apc^Min/+^ mice

**DOI:** 10.64898/2026.03.12.711485

**Authors:** Yuta Saito, Tomoko Namie, Miho Naoi, Takahiro J. Nakamura

**Affiliations:** Laboratory of Animal Physiology, School of Agriculture, Meiji University, 1-1-1 Higashimita, Tama-ku, Kawasaki, Kanagawa 214-8571, Japan

**Keywords:** Apc^min^, Clock gene, Colorectal cancer, Familial colorectal adenomatosis, β-catenin, Polyps

## Abstract

The circadian clock coordinates physiology and behavior through ∼24-h rhythms, and disruption of core clock genes has been implicated in tumorigenesis. However, the impact of *Per1*, a major clock gene, on colorectal tumor development remains unclear. Here, we investigated how *Per1* deletion influences intestinal tumorigenesis using the Apc^Min/+^ mouse model and Apc^Min/+^*Per1^-/-^*mice generated by crossing Apc^Min/+^ and *Per1^-/-^* lines (C57BL/6J background). Mice were maintained under controlled light–dark conditions, and we assessed survival, intestinal polyp burden, histopathology using Swiss-roll sections, β-catenin protein abundance (immunofluorescence and western blotting), Ctnnb1 mRNA expression (RT-qPCR), and crypt proliferation (5-bromo-2’-deoxyuridine (BrdU) immunohistochemistry).

*Per1* deletion did not significantly alter overall survival in Apc^Min/+^ mice but increased inter-individual variability. In contrast, polyp number was markedly increased by *Per1* deletion, affecting both small (<2 mm) and large (≥2 mm) polyps across intestinal segments. Histology confirmed aberrant crypt foci and polyps in both Apc^Min/+^ genotypes. β-Catenin protein levels in the whole intestine were significantly increased by *Per1* deficiency and Apc mutation (two-way ANOVA), whereas Ctnnb1 mRNA was largely unchanged across regions. BrdU-based crypt proliferation was increased by the Apc mutation but not by *Per1* deletion.

These results indicate that *Per1* loss exacerbates intestinal polyp formation and elevates β-catenin predominantly through non-transcriptional mechanisms, supporting a tumor-suppressive role of *Per1* in colorectal tumorigenesis.

## Introduction

The circadian clock receives stimuli from the external environment as cues for time information and acts as an autonomous oscillating body. It integrates and organizes time information to ensure that various behavioral and physiological processes are carried out at the appropriate times by keeping a rhythm of approximately 24 hours. In mammals, the circadian clock is genetically defined and composed of a series of genes called clock genes. When clock genes lose their function due to deficiency or other reasons, they alter various behavioral and physiological functions [1]. *Period (Per)1* is one of the core clock genes [2]. It influences the rhythm mainly through its interaction with other clock genes, while *Per2* positively regulates the expression of clock genes, and partial compensation exists between these two gene products [3].

Colorectal cancer is the third most common cancer worldwide, accounting for about 10% of all cancer cases [4]. Colorectal cancer is a general term for cancer arising from the mucosa on the surface of the colon and rectum and is a type of colorectal polyp. It is known that polyps can become malignant and become cancerous if left untreated, and the larger the polyp, the higher the risk of becoming cancerous. Among cancer suppressor genes, the Apc (adenomatous polyposis coli) gene was isolated in 1991 as the causative gene of familial colorectal adenomatosis [5]. In the process of tumorigenesis of normal colonic mucosa by inactivation of the Apc gene, the intracellular β-catenin concentration plays an important role as a switch for cell growth and differentiation. In the normal colonic epithelium, the cells near the crypt floor, including stem cells, receive Wnt signaling from stromal cells, which enhances proliferation and inhibits differentiation. After stem cells divide, daughter cells migrate upward, and Wnt stimulation from stromal cells is attenuated, resulting in rapid β-catenin degradation and cell differentiation. On the other hand, Apc gene aberrations inhibit β-catenin degradation and enhance various proliferative signals downstream of Wnt signaling. Apc mutations cause aberrant crypt foci (ACF), an early tumor precursor lesion, and adenomas [6, 7].

Apc^Min/+^ mouse (C57BL/6J- Apc^Min^/J; JAX stock #002020) is a model animal for familial adenomatous polyposis of the colon (FAP), which was created at the University of Wisconsin, USA [8]. They are widely used as mouse models of colorectal cancer to form intestinal adenomas early and spontaneously [9]. It has been reported that 100% of Apc^Min/+^ mice raised on a high-fat diet form more than 30 adenomas throughout the intestinal tract, most of which are anemic, bloody, and die by 120 days of age [8, 10]. Furthermore, homozygous individuals, Apc^Min/Min^ mice, do not survive and are embryonic lethal. It is also known that β-catenin levels are elevated in the intestine of Apc^Min/+^ mice [11], and in intestinal cells (mainly intestinal villi) of wild-type mice, β-catenin is localized in the cell cortex, while in tumors it is localized in the cell nucleus [12].

Numerous pathological studies on the clock genes that make up the circadian clock and cancer have been reported to date, many of which indicate that “clock gene defects exacerbate cancer.” Fu and colleagues showed that γ-irradiation of *Per2^-/-^* mice resulted in higher tumor development [13]. In addition, reduced *Per1* expression has been reported in several cancers, including prostate, breast, colorectal, and endometrial cancers [14, 15, 16]. Moreover, decreased *Per1* expression has been found to disrupt the balance between cell proliferation and apoptosis and promote the transformation of malignant cells [17, 18, 19].

On the other hand, some reports claim that “clock gene deletions do not exacerbate cancer”; mice lacking *Per1* and *Per2* did not suffer from spontaneous and radiation-induced cancers [20]. Mice lacking *Cryptochrome (Cry)1* and *Cry2*, other essential clock genes, were indistinguishable from wild-type mice with respect to spontaneous and radiation-induced cancer [21]. Thus, the results vary depending on the type of cancer and the type of clock gene deficiency. It is difficult to say that “clock gene deficiency aggravates cancer” in general. Although many studies have been conducted, the impact of defects in *Per1*, a major clock gene, on colorectal cancer remains unclear. In the present study, we aimed to clarify how defects in the clock gene *Per1* affect the development and progression of colorectal cancer.

## Materials and methods

### Animals

We used Apc^Min/+^ mice, a model animal for familial colorectal adenomatosis, *Per1^-/-^*mice lacking the clock gene *Per1* [3], and Apc^Min/+^*Per1^-/-^*(C57BL/6J background) mice created by crossing them. All animals were housed under controlled environmental conditions (22 ± 1°C, 50 ± 10% humidity) with a 12-h light/12-h dark cycle, with lights on at 08:00 (ZT 0) and lights off at 20:00 (ZT 12). Standard chow (MF; Oriental Yeast Co., Ltd., Tokyo, Japan) and tap water were provided ad libitum. Health checks and replenishment of food and water were performed daily between ZT 4 and ZT 8. For survival analysis, to avoid the influence of external light, mice were transferred at 8 weeks of age to a completely light-tight housing chamber (“coffin”), in which the housing conditions were identical to those described above. For all animal experiments, experimental protocols were prepared in accordance with the “Guidelines for the Appropriate Conduct of Animal Experiments” of the Science Council of Japan and the “Basic Guidelines for Animal Experiments in the Field of Physiology” of the Physiological Society of Japan, and experiments approved by the Animal Experiment Committee, Meiji University (approval number: MUIACUC2020-06) were carried out.

### Generation of Apc^Min/+^*Per1^-/-^* mice

In the F0 generation, Apc^Min/+^ mice were crossed with *Per1^-/-^*mice to obtain *Per1^-/+^* mice or Apc^Min/+^*Per1^-/+^*mice (F1 generation). Then, Apc^Min/+^*Per1^-/+^* mice obtained in the F1 generation were crossed with *Per1^-/-^* mice to obtain the F2 generation containing Apc^Min/+^*Per1^-/-^*mice, and after the F3 generation, Apc^Min/+^*Per1^-/-^* mice were crossed with *Per1^-/-^* mice to obtain Apc^Min/+^*Per1^-/-^* mice. *Per1^-/-^* mice were obtained by crossing Apc^Min/+^*Per1^-/-^* mice with *Per1^-/-^* mice. All Apc^Min/+^ mice used in the mating were male mice to avoid the death of the mother mice or abandonment of nursing during the neonatal period of the pups. In the experiments, mice were used so that male: female = 1:1. Because Apc^Min/+^ and *Per1^-/+^* mice have been bred on a C57BL/6J background for more than 20 generations each, C57BL/6J mice that are home-bred as a separate colony in our laboratory were used as wild-type (WT) mice in the experiment.

### Recording the number of survival days

Survival days of Apc^Min/+^ and Apc^Min/+^*Per1^-/-^* mice were recorded. The weight of each individual was measured and recorded weekly with an animal balance (Shimadzu Corporation, Kyoto, Japan) starting at 8 weeks of age. Health checks were performed daily between ZT 4 and ZT 8. For humane endpoints, mice were placed under watchful waiting if they lost 10% or more of their body weight at 8 weeks of age and were euthanized by cervical dislocation if they lost 20% or more of their body weight.

### Polyp count measurement

We measured the number of polyps in Apc^Min/+^ and Apc^Min/+^*Per1^-/-^*mice, respectively. For the present study, 150-day-old mice were used, which were close to death due to advanced polyp formation. Mice were euthanized by cervical dislocation, and the intestinal tract from just after the stomach to the anus was removed; after washing with phosphate-buffered saline (PBS), a longitudinal incision was made. After the incision, the contents of the intestinal tract were removed with tweezers, and the intestinal tract was rewashed with PBS. The removed intestinal tract was defined as the small intestine from just after the stomach to just before the cecum and the colon from just after the cecum to the anus. The small intestine was further divided into four equal parts, the duodenum, jejunum 1, jejunum 2, and ileum from the top, and the intestinal tissue was placed on a 2 mm square Petri dish. Polyps less than 2 mm in diameter were defined as small polyps, and polyps greater than 2 mm as large polyps.

### Histological evaluation

Swiss roll is a useful analysis method that allows unbroken observation of the entire intestinal tract on a single slide [22]. In the present study, 150-day-old mice were used. Intestinal rolls were prepared by wrapping the intestinal tissue around a toothpick while fixing it with 4% paraformaldehyde (PFA) and securing it with a gusset needle. They were then immersion-fixed in 4% PFA, and then they were immersed in 30% sucrose solution the next day. The 50 µm thick intestinal roll sections were prepared using a cryostat (Thermo Fisher Scientific, Massachusetts, USA) and mounted on glass slides (Matsunami Glass, Osaka, Japan). After sectioning, each intestine was stained with hematoxylin and eosin (H&E) stain for histological evaluation.

In addition, immunofluorescence staining was performed (primary antibody: Beta Catenin Polyclonal antibody, Proteintech Japan, Tokyo, Japan; secondary antibody: Goat anti-Rabbit IgG (H+L) Highly Cross Adsorbed Secondary Antibody, Alexa Fluor 488, Thermo Fisher Scientific). The nuclei were contrast-stained blue with DAPI to confirm the extent of the intestine. After staining, the sections were sealed in 1 x PBS and cover glass. The sealed sections were observed using a fluorescence microscope (ZEISS, Oberkochen, Germany). Specificity was confirmed by the removal of the primary antibody.

### Western blotting

Western blotting method was used as previously reported [23]. Briefly, using 150-day-old mice, as in the polyp-count analysis, the intestine was collected between ZT 6 and ZT 8. The excised intestine was divided as described before, and normal tissue excluding tumor-like lesions was harvested. After homogenization, it was centrifuged at 12,000 g for 15 min (Kubota, Osaka, Japan). The supernatant was collected and the total protein concentration was determined by the Bicinchoninic Acid (BCA) method. The protein bands were detected using an ImageQuant LAS 4000 Mini Luminescent Image Analyzer (GE Healthcare, Chicago, IL, United States) and ECL Prime Western Blotting Detection Kit (GE Healthcare). Actin (Beta Actin Polyclonal antibody, Proteintec Japan) was used as a loading control, and the final β-catenin expression level is the optical density of the β-catenin band simultaneously flowing divided by the optical density of the actin band obtained by western blotting. ImageJ (National Institutes of Health, Maryland, USA) image processing software was used to process and analyze the images.

### Real-time quantitative PCR (RT-qPCR)

In this experiment, we used 120-day-old mice, which allowed us to secure a sufficient sample size, and collected intestinal tissues at ZT 8. The excised intestine was subdivided as described before, and normal tissue excluding tumor-like lesions was harvested.

Total RNA was extracted from each intestinal segment using TRIzol® Reagent (Thermo Fisher Scientific), and cDNA was synthesized from total RNA using ReverTra Ace® qPCR RT Master Mix (TOYOBO, Osaka, Japan). RNA extraction and cDNA synthesis were performed according to the manufacturers’ protocols.

qPCR was performed using a StepOnePlus Real-Time PCR System (Thermo Fisher Scientific) with THUNDERBIRD™ Probe qPCR Mix. The thermal cycling conditions were as follows: 95°C for 10 min, followed by 40 cycles of 95°C for 5 s and 60°C for 15 s. A pooled cDNA (equal amounts from all samples) was serially diluted to generate a standard curve for relative quantification. Values were normalized to Rplp0 and expressed relative to the WT mean (=1). qPCR procedures were conducted in accordance with the manufacturers’ protocols. Probes were obtained from PrimeTime Assays (Integrated DNA Technologies, IA, USA) and were designed to span exon–exon junctions [Rplp0 (Mm.PT.58.438942059), Ctnnb1 (Mm.PT.58.12501105)].

### Cell counting by immunohistochemistry

In this experiment, 120–140-day-old mice were used. BrdU (Sigma-Aldrich, St. Louis, MO, USA) was prepared at 10 mg/mL in PBS and administered intraperitoneally at 100 mg/kg 1.5 h prior to intestinal collection. Mice were euthanized at ZT 8, and the entire intestine from immediately distal to the stomach to the anus was excised. Intestinal Swiss-roll sections were prepared as described above, and the section thickness was set to 10 μm. Sections were subjected to BrdU immunohistochemistry to detect BrdU-positive nuclei (VECTASTAIN® ABC-HRP Kit; Vector Laboratories, CA, USA; ImmPACT® DAB Substrate, Peroxidase (HRP); Vector Laboratories; primary antibody: anti-BrdU antibody; Abcam, Cambridge, UK), followed by hematoxylin counterstaining. Immunohistochemistry was performed according to the manufacturers’ recommended protocols. Images were acquired across the entire intestinal region and analyzed in a manner blinded to animal ID.

Quantification of the total number of cells and proliferating cells (DAB-positive) in intestinal crypts was performed using Python. For each image, a crypt-containing region was manually cropped. To account for inter-individual variation in staining intensity, representative images with different DAB intensities were selected from each animal, and parameters were adjusted to minimize the difference between automated and manual counts.

The analysis code and parameters are provided in a public repository.

### Statistical Analysis

Survival days and polyp count for Apc^Min/+^ mice and Apc^Min/+^*Per1^-/-^* mice are shown as mean ± standard deviation (SD), and Welch’s t-test was used to compare the two groups. Survival rates of Apc^Min/+^ mice and Apc^Min/+^*Per1^-/-^* mice were calculated by the Kaplan-Meier method, and survival curves were generated. A log-rank test was performed on the generated survival curves. β-Catenin protein expression levels, Ctnnb1 mRNA expression levels, and the proportion of proliferating cells are presented as mean ± standard error of the mean (SEM). Comparisons among the four groups were performed using two-way analysis of variance (two-way ANOVA), with *Per1* deficiency and the Apc mutation as the two factors. *P* values for each test were performed at 5%, with *P* < 0.05 indicating a significant difference.

## Results

### Effect of *Per1* deletion on survival days and survival rate of Apc^Min/+^ mice

The number of survival days of Apc^Min/+^ and Apc^Min/+^*Per1^-/-^*mice were recorded to determine the effect of *Per1* deletion on the number of survival days and survival rate of Apc^Min/+^ mice. The number of survival days was 158.16 ± 26.51 (mean ± SD, same as below) days for Apc^Min/+^ mice (n = 68) and 168.67 ± 58.70 days for Apc^Min/+^*Per1^-/-^* mice (n = 49), with no significant differences (Figure 1A; *P* = 0.25, Welch’s t-test).

**Figure 1:**
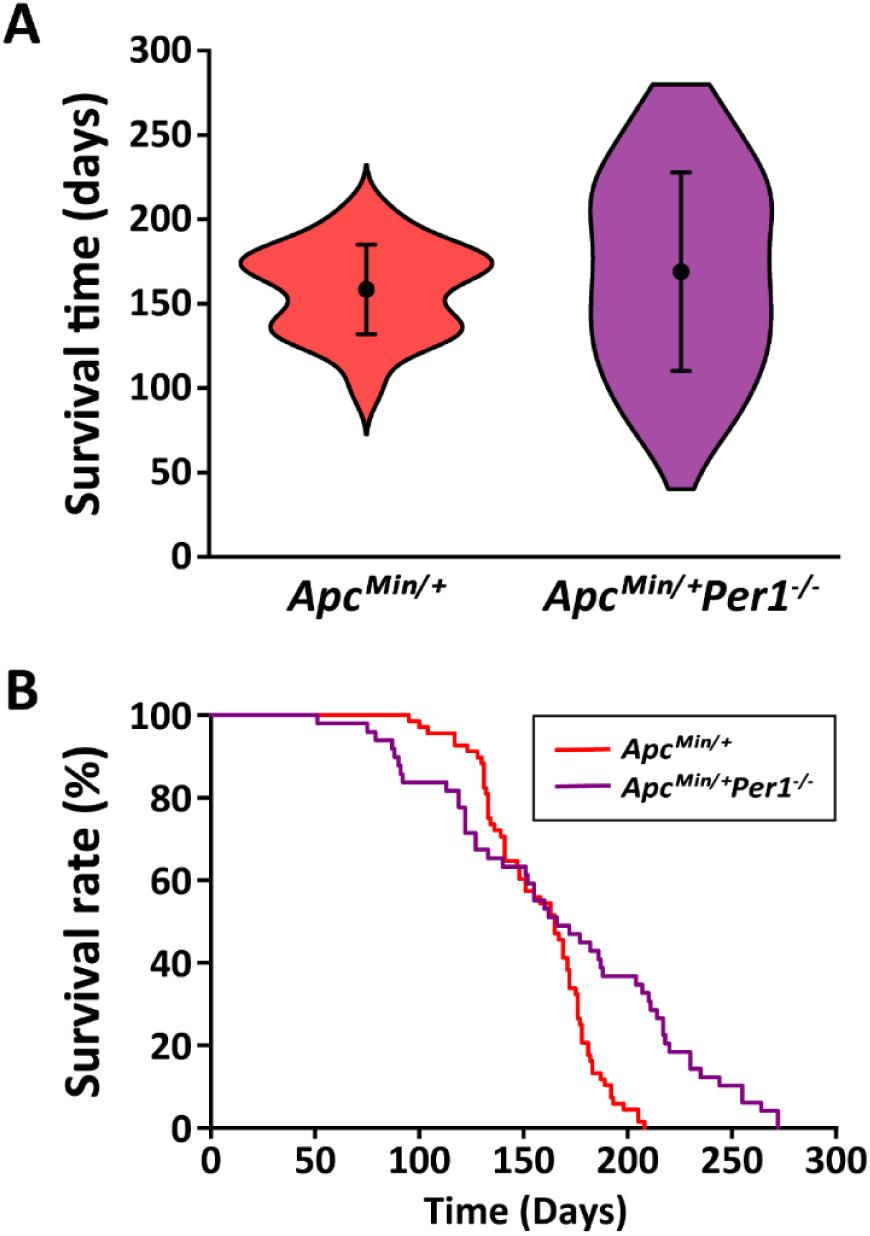
Survival days and survival rate of Apc^Min/+^ and Apc^Min/+^*Per1^-/-^* mice. (A) Violin plots of survival days for Apc^Min/+^ and Apc^Min/+^*Per1^-/-^* mice are shown. (B) Kaplan-Meier survival curves for Apc^Min/+^ and Apc^Min/+^*Per1^-/-^* mice are shown.

From the obtained data on the number of survival days, the survival rate of Apc^Min/+^ and Apc^Min/+^*Per1^-/-^*mice was calculated using the Kaplan-Meier method, and survival curves were generated (Figure 1B). Although the survival rate of Apc^Min/+^*Per1^-/-^*mice was lower than that of Apc^Min/+^ mice up to 140 days of age when the survival rate reached 60%, the difference disappeared when the survival rate fell below 60% and reversed below 50%. Log-rank test showed no significant difference (*P* = 0.09, Log-rank test).

### Effect of *Per1* deletion on the number of polyps in the intestinal tract of Apc^Min/+^ mice

We measured the number of polyps in Apc^Min/+^ and Apc^Min/+^*Per1^-/-^* mice to determine the effect of *Per1* deficiency on the number of polyps in Apc^Min/+^ mice. The number of polyps was 26.86 ± 15.42 (mean ± SD, same as below) in Apc^Min/+^ mice (n = 21) and 54.80 ± 23.41 in Apc^Min/+^*Per1^-/-^*mice (n = 20). The number of polyps in Apc^Min/+^*Per1^-/-^* mice was significantly higher than that of Apc^Min/+^ mice (Figure 2A; *P* < 0.05, Welch’s t-test).

**Figure 2:**
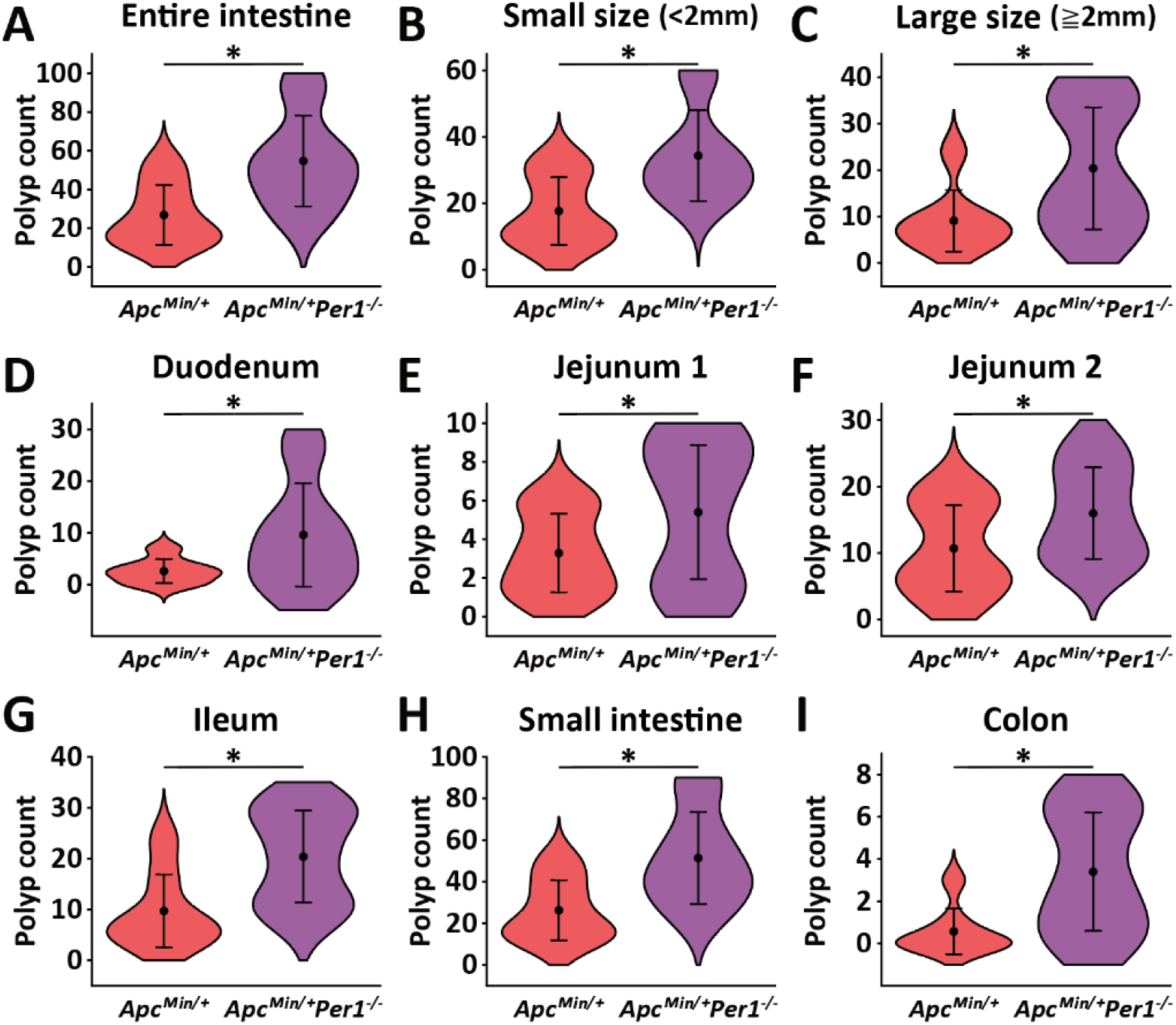
Polyp counts of 150-day-old Apc^Min/+^ and Apc^Min/+^*Per1^-/-^* mice. (A) Violin plots of polyp counts across the intestine of Apc^Min/+^ and Apc^Min/+^*Per1^-/-^* mice are shown. (B, C) Violin plots of polyp count of small polyps (B, < 2 mm diameter) and large polyps (C, > 2 mm diameter) in Apc^Min/+^ and Apc^Min/+^*Per1^-/-^*mice are shown. (D-I) Violin plots of polyp counts in duodenum (D), jejunum 1 (E), jejunum 2 (F), ileum (G), small intestine (H, duodenum to ileum), and colon (I) of Apc^Min/+^ and Apc^Min/+^*Per1^-/-^*mice are shown. *: *P* < 0.05, Welch’s t-test.

Next, we divided the polyp counts into small and large polyps and compared the number of polyps in each. The number of small polyps was 17.71 ± 10.15 for Apc^Min/+^ mice and 34.40 ± 13.67 for Apc^Min/*+*^*Per1^-/-^* mice. The number of large polyps was 9.14 ± 6.63 for Apc^Min/+^ mice and 20.40 ± 13.15 for Apc^Min/+^*Per1^-/-^* mice. For both small and large polyps, Apc^Min/+^*Per1^-/-^* mice had more polyps than Apc^Min/+^ mice (Figure 2B, C; small polyps: *P* < 0.05, Welch’s t-test; large polyps: *P* < 0.05, Welch’s t-test).

The intestines were further classified into duodenum, jejunum 1, jejunum 2, ileum, small intestine (duodenum to ileum), and colon, and the number of polyps was compared in each area. In the duodenum, Apc^Min/+^ mice had 2.57 ± 2.31 polyps and Apc^Min/+^*Per1^-/-^* mice had 9.60 ± 10.02 polyps; in the jejunum 1, Apc^Min/+^ mice had 3.29 ± 2.03 polyps and Apc^Min/+^*Per1^-/-^* mice had 5.40 ± 3.47 polyps; in the jejunum 2, Apc^Min/+^ mice had 10.71 ± 6.54 polyps and Apc^Min/+^*Per1^-/-^*mice had 16.00 ± 6.90 polyps; in the ileum, Apc^Min/+^ mice had 9.71 ± 7.18 polyps and Apc^Min/+^*Per1^-/-^*mice had 20.40 ± 9.03 polyps; in the small intestine, Apc^Min/+^ mice had 26.29 ± 14.49 polyps and Apc^Min/*+*^*Per1^-/-^*mice had 51.40 ± 22.24 polyps; and in the colon, Apc^Min/+^ mice had 0.57 ± 1.08 polyps and Apc^Min/+^*Per1^-/-^*mice had 3.40 ± 2.80 polyps. In all areas, the numbers of polyps were significantly higher in Apc^Min/+^*Per1^-/-^* mice than those in Apc^Min/+^ mice (Figure 2D-I; *P* < 0.05, Welch’s t-test).

### Effect of *Per1* deletion on intestinal tissue of Apc^Min/+^ mice

H&E staining was performed after intestinal roll sections were prepared, and the stained images of each section are shown in Figure 3 to compare the intestinal tissue structures of WT, *Per1^-/-^*, Apc^Min/+^, and Apc^Min/+^*Per1^-/-^* mice. The sections of WT and *Per1^-/-^* mice showed normal intestinal structure, with uniform crypts and villi observed. On the other hand, intestinal roll sections from Apc^Min/+^ and Apc^Min/+^*Per1^-/-^* mice showed abnormal crypt foci (ACF). Several polyps were also observed. Furthermore, immunofluorescent staining for β-catenin in intestinal roll sections confirmed the expression of β-catenin in the intestinal villi in sections from *Per1^-/-^*, Apc^Min/+^, and Apc^Min/+^*Per1^-/-^*mice. We also observed β-catenin expression near the crypts in sections from Apc^Min/+^*Per1^-/-^* mice (Figure 4).

**Figure 3:**
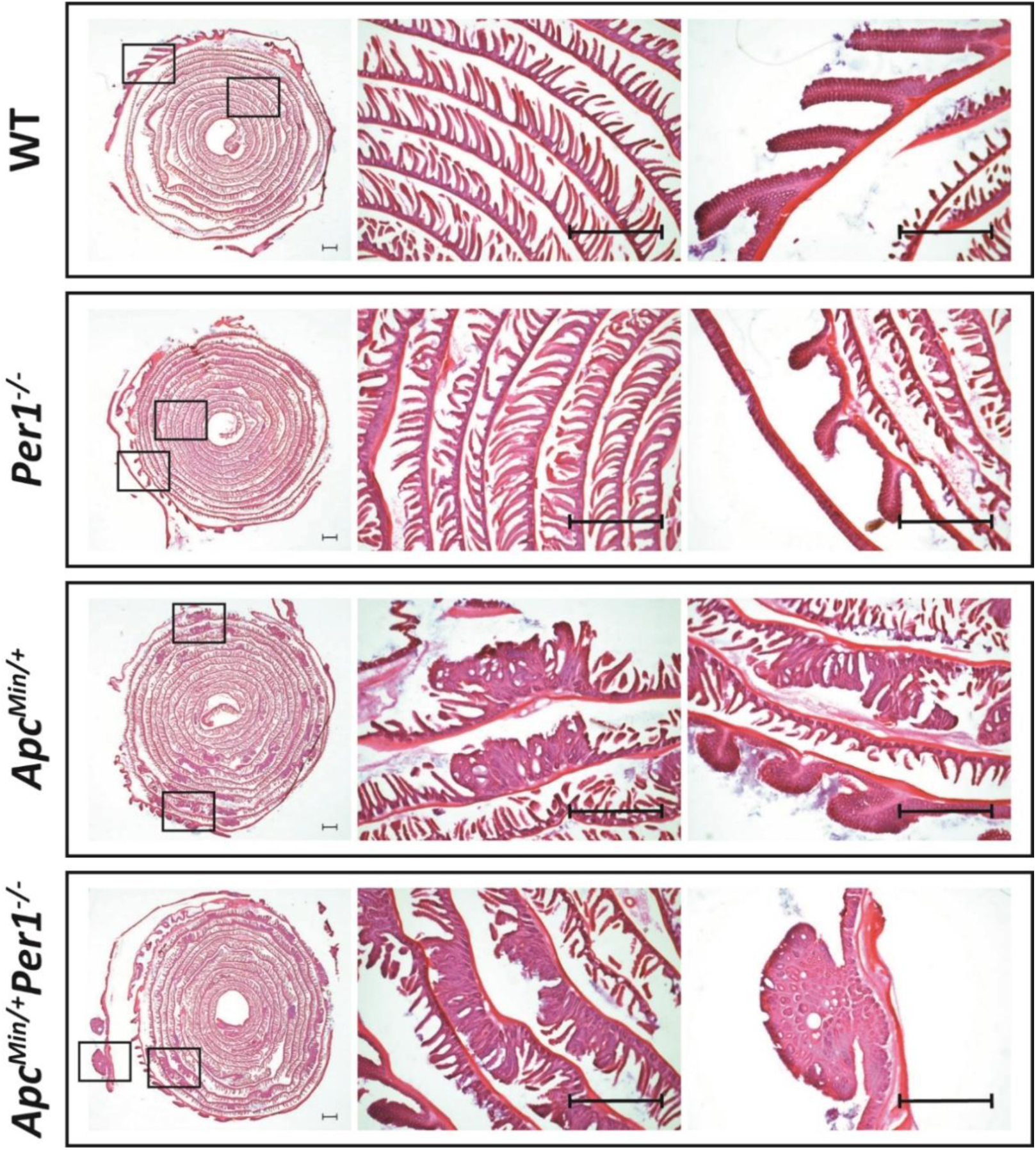
H&E staining images in intestinal roll sections. Bright field Images of Intestinal roll sections of WT, *Per1^-/-^*, Apc^Min/+^, and Apc^Min/+^*Per1^-/-^* mice after H&E staining are shown. The left image is a x1 stained image, and the center and right images are x4 stained images. Black squares in the left image indicate the location of the center and right images. Black scale bars indicate 1 mm.

**Figure 4:**
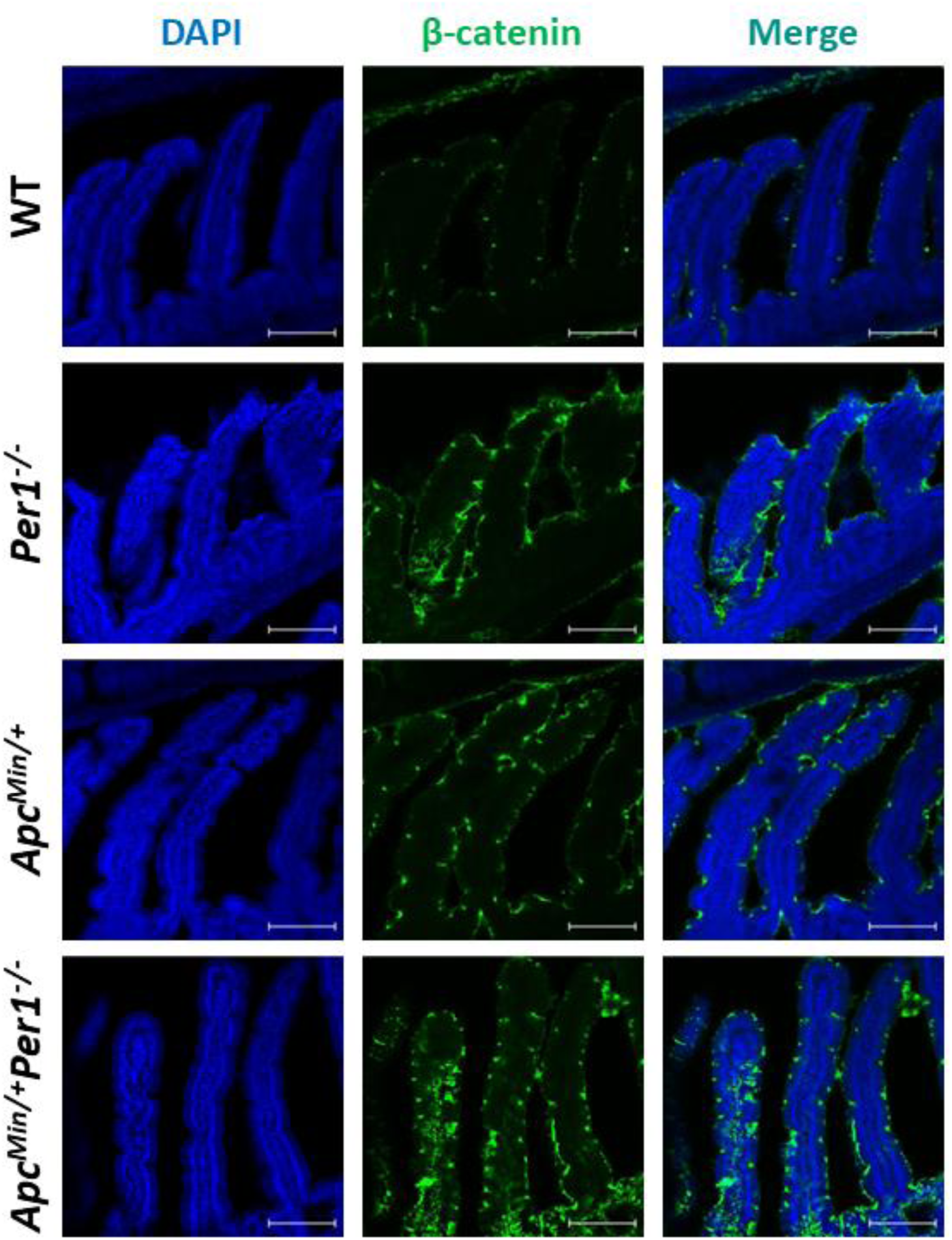
Immunofluorescence staining images of β-catenin expression in intestinal roll sections. Fluorescence images of Intestinal roll sections of WT, *Per1^-/-^*, Apc^Min/+^, and Apc^Min/+^*Per1^-/-^*mice after immunofluorescence staining are shown. The left images show DAPI, the middle image shows β-catenin, and the right images show their merged images. They are x10 stained images. The white scale bars indicate 100 µm.

### Effect of *Per1* deletion on intestinal β-catenin expression in mice

To ascertain whether the polyp development caused by Per1 deficiency is mediated by β-catenin, we measured the expression level of β-catenin in the intestine of WT, *Per1^-/-^*, Apc^Min/+^, and Apc^Min/+^*Per1^-/-^* mice (n = 3 each) by western blotting (Figure 5A). The results showed that the overall intestinal expression levels were 1.00 ± 0.32 (mean ± SEM, same as below) in WT mice, 8.97 ± 3.61 for *Per1^-/-^* mice, 43.27 ± 8.71 in Apc^Min/+^ mice, and 50.73 ± 3.18 in Apc^Min/+^*Per1^-/-^*mice. No significant differences were found in the interaction, but significant differences were found in the *Per1* deficiency and Apc mutation factors (Figure 5B; *Per1* deficiency factor: *P* < 0.05; Apc mutation factor: *P* < 0.001; interaction: *P* = 0.93, Two-way ANOVA).

**Figure 5:**
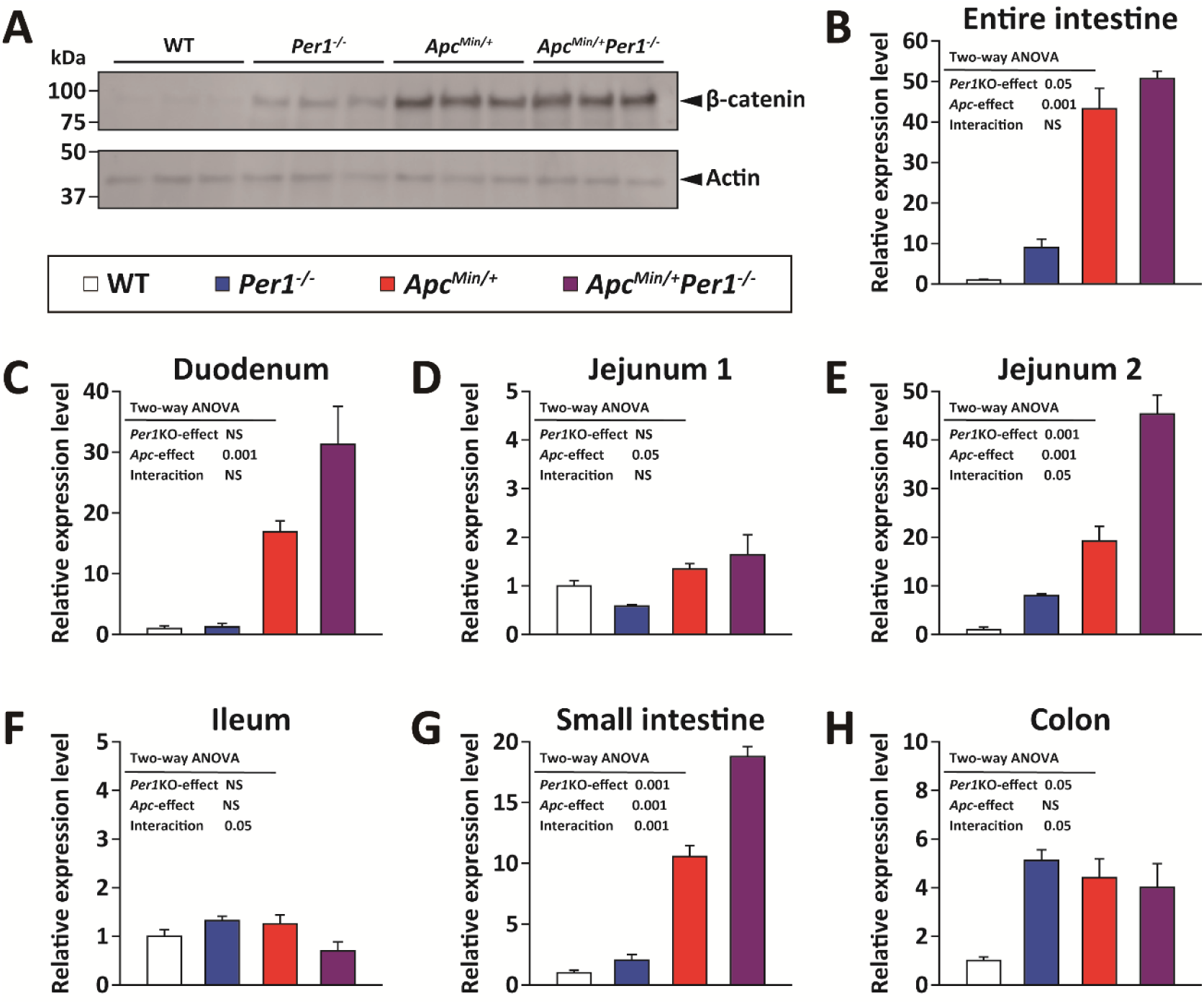
β-catenin expressions in the intestine of WT, *Per1^-/-^*, Apc^Min/+^, and Apc^Min/+^*Per1^-/-^*mice. (A) Western blotting images in the whole intestine of WT, *Per1^-/-^*, Apc^Min/+^, and Apc^Min/+^*Per1^-/-^*mice are shown. The upper panel shows β-catenin and the lower panel shows actin. (B) Quantitative results of western blotting of β-catenin in the whole intestine of WT, *Per1^-/-^*, Apc^Min/+^, and Apc^Min/+^*Per1^-/-^* mice (relative values with WT mice as 1) are shown. (C-H) Quantitative results of western blotting of β-catenin in duodenum (C), jejunum 1 (D), jejunum 2 (E), ileum (F), small intestine (G, duodenum to ileum) and colon (H) of WT mice, *Per1^-/-^* mice, Apc^Min/+^ mice, Apc^Min/+^*Per1^-/-^* mice (relative values with WT mice as 1) are shown.

In the duodenum, levels were 1.00 ± 0.72 in WT mice, 1.27 ± 0.94 in *Per1^-/-^* mice, 16.90 ± 3.08 in Apc^Min/+^ mice, and 31.29 ± 10.80 Apc^Min/+^*Per1^-/-^*mice, no significant differences were found in the *Per1* deficiency factor and interaction, but significant differences were found in the Apc mutation factor (Figure 5C; *Per1* deficiency factor: *P* = 0.06; Apc mutation factor: *P* < 0.001; interaction: *P* = 0.06, Two-way ANOVA).

In the jejunum 1, levels were 1.00 ± 0.19 in WT mice, 0.58 ± 0.06 in *Per1^-/-^* mice, 1.34 ± 0.19 in Apc^Min/+^ mice, and 1.64 ± 0.71 in Apc^Min/+^*Per1^-/-^*mice, no significant differences were found in the *Per1* deficiency factor and interaction, but significant differences were found in the Apc mutation factor (Figure 5D; *Per1* deficiency factor: *P* = 0.79; Apc mutation factor: *P* < 0.05; interaction: *P* = 0.14, Two-way ANOVA).

In the jejunum 2, levels were 1.00 ± 0.90 in WT mice, 8.01 ± 0.61 in *Per1^-/-^* mice, 19.24 ± 5.17 in Apc^Min/+^ mice, and 45.35 ± 6.73 in Apc^Min/+^*Per1^-/-^*mice, the factors of *Per1* deficiency, Apc mutation, and interaction showed significantly different (Figure 5E; *Per1* deficiency factor: *P* < 0.001; Apc mutation factor: *P* < 0.001; interaction: *P* < 0.05, Two-way ANOVA).

In the ileum, levels were 1.00 ± 0.24 in WT mice, 1.32 ± 0.15 in *Per1^-/-^* mice, 1.26 ± 0.32 in Apc^Min/+^ mice, and 0.70 ± 0.32 in Apc^Min/+^*Per1^-/-^*mice, no significant differences were found in the *Per1* deficiency and Apc mutation factors, but significant differences were found in the interaction factor (Figure 5F; *Per1* deficiency factor: *P* = 0.47; Apc mutation factor: *P* = 0.27; interaction: *P* < 0.05, Two-way ANOVA).

In the small intestine, levels were 1.00 ± 0.38 in WT mice, 2.03 ± 0.80 in *Per1^-/-^*mice, 10.58 ± 1.53 in Apc^Min/+^ mice, and 18.79 ± 1.42 in Apc^Min/+^*Per1^-/-^*mice, the factors of *Per1* deficiency, Apc mutation, and interaction showed significantly different (Figure 5G; *Per1* deficiency factor: *P* < 0.001; Apc mutation factor: *P* < 0.001; interaction: *P* < 0.001, Two-way ANOVA).

In the colon, levels were 1.00 ± 0.26 in WT mice, 5.11 ± 0.78 in *Per1^-/-^* mice, 4.42 ± 1.33 in Apc^Min/+^ mice, and 4.01 ± 1.70 in Apc^Min/+^*Per1^-/-^*mice, no significant differences were found in the Apc mutation factor, but significant differences were found in the *Per1* deficiency factor and interaction (Figure 5H; *Per1* deficiency factor: *P* < 0.05; Apc mutation factor: *P* = 0.12; interaction: *P* < 0.05, Two-way ANOVA).

### Effect of *Per1* deficiency on Ctnnb1 mRNA expression in the mouse intestine

To determine whether the *Per1* deficiency–induced increase in β-catenin protein expression is associated with transcriptional changes, we measured Ctnnb1 (β-catenin) mRNA levels in intestinal tissues from WT, *Per1^-/-^*, Apc^Min/+^, and Apc^Min/+^ *Per1*^-/-^ mice (n = 3 each) by RT-qPCR (Fig. 6). In the whole intestine, the relative Ctnnb1 mRNA level was 1.00 ± 0.22 in WT mice, 1.14 ± 0.45 in *Per1*^-/-^ mice, 1.31 ± 0.13 in Apc^Min/+^ mice, and 1.31 ± 0.12 in Apc^Min/+^ *Per1*^-/-^ mice (mean ± SEM; same hereafter). No significant main effects of *Per1* deficiency or Apc mutation, nor their interaction, were detected (Fig. 6A; *Per1* deficiency: *P* = 0.80; Apc mutation: *P* = 0.10; interaction: *P* = 0.82; two-way ANOVA).

**Figure 6.**
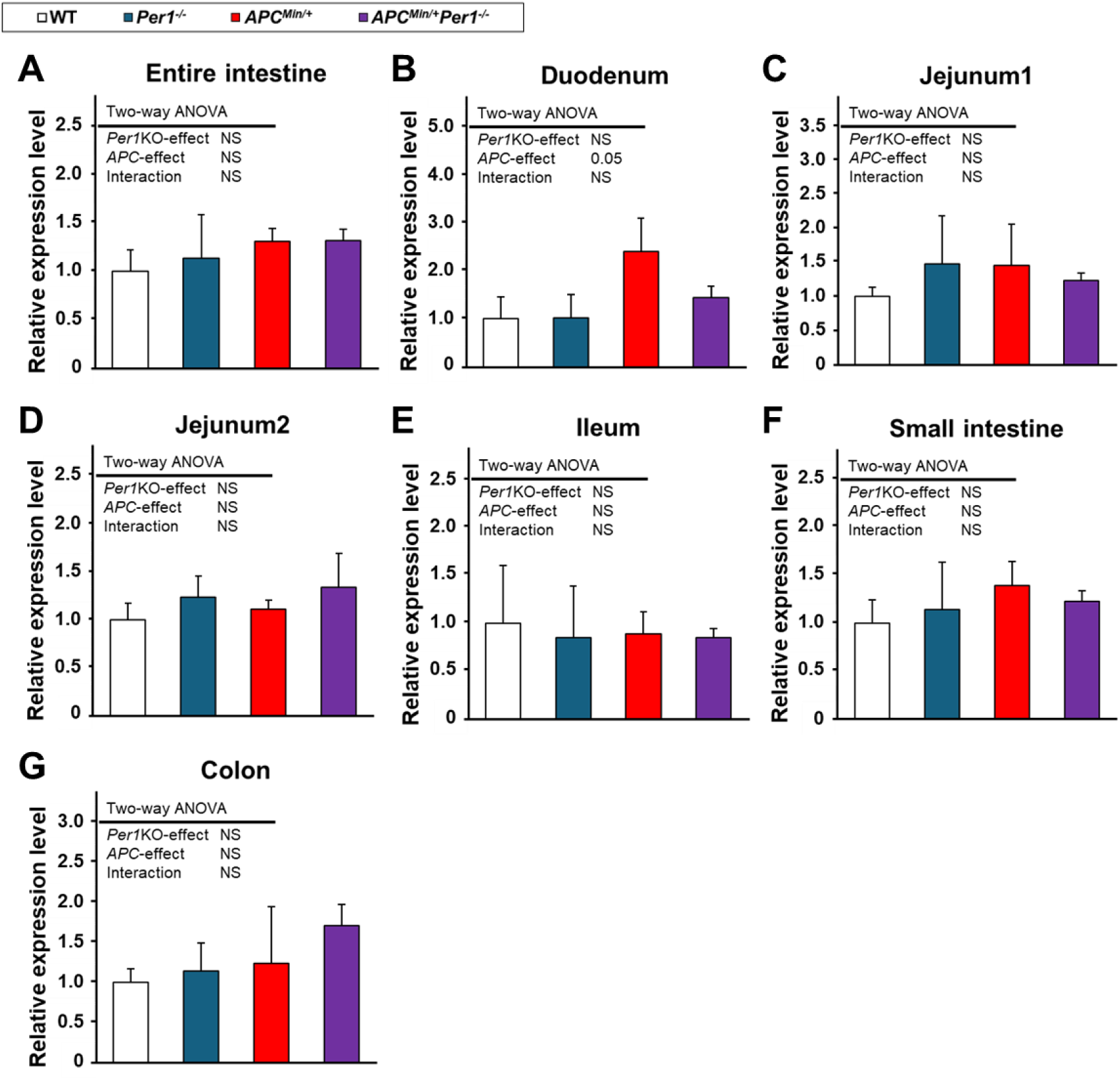
Ctnnb1 mRNA expression in the intestine of WT, *Per1^-/-^*, Apc^Min/+^, and Apc^Min/+^ *Per1^-/-^ mice.* (A) RT-qPCR quantification of Ctnnb1 in the whole intestine of WT, *Per1^-/-^*, Apc^Min/+^, and Apc^Min/+^*Per1^-/-^* mice, shown as relative values normalized to WT (WT = 1). The y-axis indicates relative β-catenin mRNA expression. (B–G) RT-qPCR quantification of Ctnnb1 in the duodenum (B), jejunum 1 (C), jejunum 2 (D), ileum (E), small intestine (F; duodenum to ileum), and colon (G) of WT, *Per1^-/-^*, Apc^Min/+^, and Apc^Min/+^*Per1^-/-^* mice, shown as relative values normalized to WT (WT = 1). The y-axis indicates relative β-catenin mRNA expression. P < 0.05 (two-way ANOVA).

In the duodenum, levels were 1.00 ± 0.46 in WT mice, 1.02 ± 0.49 in *Per1*^-/-^ mice, 2.40 ± 0.70 in Apc^Min/+^ mice, and 1.45 ± 0.24 in Apc^Min/+^ *Per1*^-/-^ mice. While neither the *Per1* deficiency main effect nor the interaction was significant, the Apc mutation main effect was significant (Fig. 6B; *Per1* deficiency: *P* > 0.99; Apc mutation: *P* < 0.05; interaction: *P* = 0.42; two-way ANOVA).

In jejunum 1, levels were 1.00 ± 0.14 in WT mice, 1.48 ± 0.71 in *Per1*^-/-^ mice, 1.46 ± 0.61 in Apc^Min/+^ mice, and 1.23 ± 0.12 in Apc^Min/+^ *Per1*^-/-^ mice, with no significant main effects or interaction (Fig. 6C; *Per1* deficiency: *P* = 0.33; Apc mutation: *P* = 0.28; interaction: *P* = 0.32; two-way ANOVA).

In jejunum 2, levels were 1.00 ± 0.18 in WT mice, 1.24 ± 0.22 in *Per1*^-/-^ mice, 1.11 ± 0.10 in Apc^Min/+^ mice, and 1.34 ± 0.36 in Apc^Min/+^ *Per1*^-/-^ mice, with no significant main effects or interaction (Fig. 6D; *Per1* deficiency: *P* = 0.26; Apc mutation: *P* = 0.45; interaction: *P* = 0.86; two-way ANOVA).

In the ileum, levels were 1.00 ± 0.60 in WT mice, 0.85 ± 0.54 in *Per1*^-/-^ mice, 0.89 ± 0.23 in Apc^Min/+^ mice, and 0.85 ± 0.10 in Apc^Min/+^ *Per1*^-/-^ mice, with no significant main effects or interaction (Fig. 6E; *Per1* deficiency: *P* = 0.74; Apc mutation: *P* = 0.95; interaction: *P* = 0.78; two-way ANOVA).

In the small intestine (combined), levels were 1.00 ± 0.24 in WT mice, 1.14 ± 0.49 in *Per1*^-/-^mice, 1.39 ± 0.25 in Apc^Min/+^ mice, and 1.23 ± 0.11 in Apc^Min/+^*Per1*^-/-^ mice, with no significant main effects or interaction (Fig. 6F; *Per1* deficiency: *P* = 0.84; Apc mutation: *P* = 0.12; interaction: *P* = 0.63; two-way ANOVA).

In the colon, levels were 1.00 ± 0.17 in WT mice, 1.14 ± 0.35 in *Per1*^-/-^ mice, 1.24 ± 0.71 in Apc^Min/+^ mice, and 1.71 ± 0.26 in Apc^Min/+^ *Per1*^-/-^ mice, with no significant main effects or interaction (Fig. 6G; *Per1* deficiency: *P* = 0.64; Apc mutation: *P* = 0.94; interaction: *P* = 0.54; two-way ANOVA).

### Effect of *Per1* deficiency on the number of proliferating cells in mouse intestinal crypts

To examine whether the *Per1* deficiency–induced increase in β-catenin protein expression affects IEC proliferation, we evaluated the proportion of proliferating cells in intestinal crypts from WT, *Per1*^-/-^, Apc^Min/+^, and Apc^Min/+^ *Per1*^-/-^ mice (n = 3 each) by immunohistochemistry (Fig. 7A). The proportion of proliferating cells was 0.193 ± 0.027 in WT mice, 0.223 ± 0.012 in *Per1*^-/-^ mice, 0.305 ± 0.018 in Apc^Min/+^ mice, and 0.335 ± 0.045 in Apc^Min/+^*Per1^-/-^*mice (mean ± SEM). While neither the *Per1* deficiency main effect nor the interaction was significant, the Apc mutation main effect was significant (Fig. 7B; *Per1* deficiency: *P* = 0.11; Apc mutation: *P* < 0.001; interaction: *P* > 0.99; two-way ANOVA).

**Figure 7.**
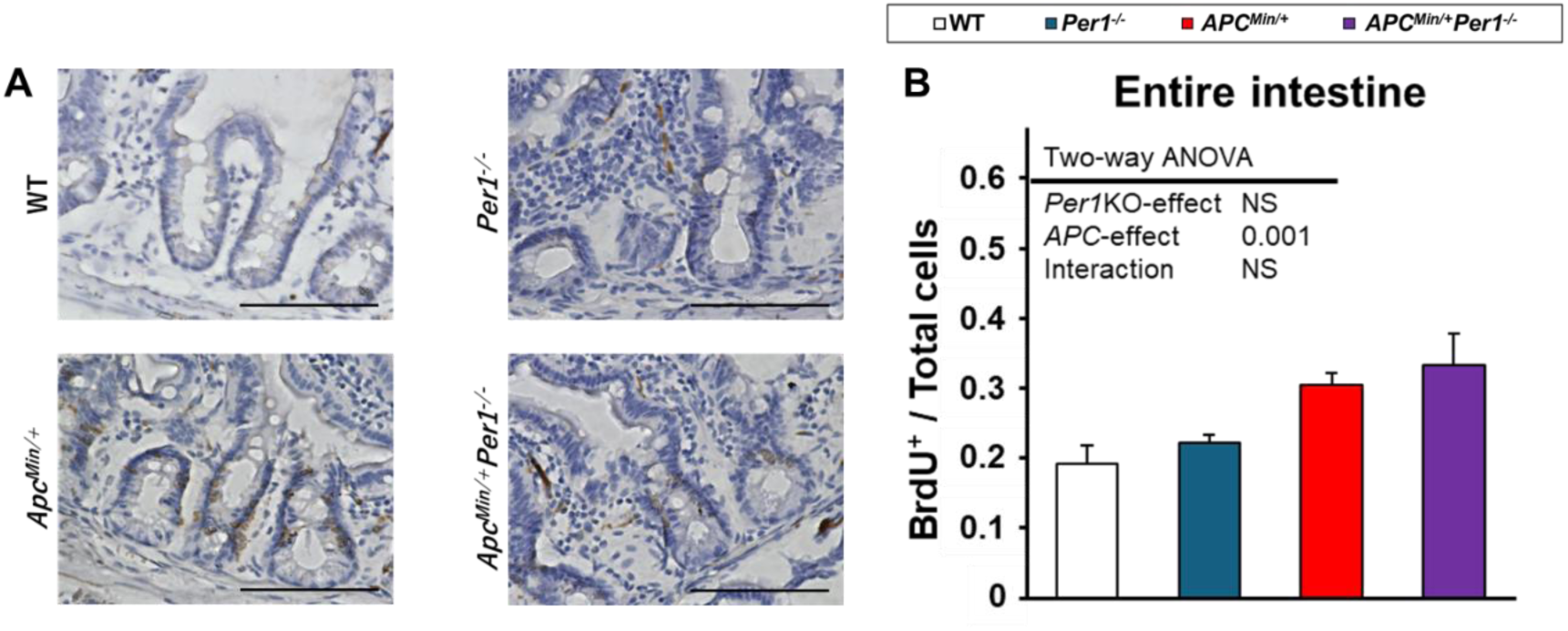
Proportion of proliferating cells in intestinal crypts of WT, *Per1^-/-^*, Apc^Min/+^, and Apc^Min/+^ *Per1^-/-^ mice*. (A) Representative immunohistochemical images acquired at ×40 from WT, *Per1^-/-^*, Apc^Min/+^, and Apc^Min/+^*Per1^-/-^*mice. Scale bar, 100 µm. (B) Proportion of proliferating cells in intestinal crypts from WT, *Per1^-/-^*, Apc^Min/+^, and Apc^Min/+^*Per1^-/-^*mice. The y-axis indicates the ratio of BrdU-positive cells to hematoxylin-positive cells within intestinal crypts (*P* < 0.001; two-way ANOVA).

## Discussion

We investigated the effects of *Per1* loss on colorectal cancer initiation and progression by analyzing survival, polyp number, intestinal histology, crypt cell proliferation, and β-catenin protein and mRNA expression. *Per1* deletion did not significantly affect survival in Apc^Min/+^ mice, but significantly increased polyp number. Aberrant crypt foci and polyps were observed in both Apc^Min/+^ and Apc^Min/+^*Per1^-/-^* mice. β-catenin was detected in Per1-/-, Apc^Min/+^, and Apc^Min/+^*Per1^-/-^*intestinal sections, and its expression in the whole intestine was significantly increased by both *Per1* deletion and the Apc mutation, despite regional variation. By contrast, *Per1* deficiency did not significantly affect Ctnnb1 mRNA expression or crypt cell proliferation.No statistically significant differences were observed in survival days or survival rate between Apc^Min/+^ and Apc^Min/+^*Per1^-/-^*mice. However, Apc^Min/+^*Per1^-/-^* mice had longer mean survival days, a larger standard deviation and a more gradual slope of the survival curve than Apc^Min/+^ mice. In other words, loss of *Per1* tends to increase inter-individual variation in survival on the Apc^Min/+^ background. The mean survival days for Apc^Min/+^ mice is reported to be 120 days [8, 10], however, it was 158 days in the present study. Given that Apc^Min/+^ survival varies substantially across individuals, the prolonged survival may reflect the use of a line repeatedly bred in our laboratory for more than 20 generations. Although Apc^Min/+^ mice reportedly die around 120 days due to anemia and bloody stools associated with intestinal narrowing and bleeding from polyps [8, 10], the exact cause of death was difficult to determine in our study, and the reason why Apc^Min/+^*Per1^-/-^* mice survived longer is unknown.

Polyp counts in Apc^Min/+^ and Apc^Min/+^*Per1^-/-^* mice revealed that *Per1* deficiency increased polyp counts. A study focusing on the clock gene *Bmal1* reported that Apc^+/-^*Bmal1^-/-^* mice had increased numbers of small intestinal polyps compared to Apc^+/-^ mice [24]. Furthermore, a study focusing on *Per2* revealed that Apc^Min/+^*Per2^m/m^*mice with mutations in *Per2* had twice as many small and large intestinal polyps as Apc^Min/+^ mice, with more severe anemia and splenomegaly [25]. In the present study, the small intestine was divided into four equal areas or measured separately by size, and in all areas and in all measurements by size, the number of polyps was significantly higher in Apc^Min/+^*Per1^-/-^* mice than Apc^Min/+^ mice. The study focusing on *Per2* found no significant differences in duodenal and large polyp counts [25]. Therefore, a lack of *Per1* may have a more significant impact on polyp count than that of *Per2*.

H&E staining of intestinal roll sections from each mouse revealed that sections from WT and *Per1^-/-^* mice showed normal intestinal structure with uniform crypts and villi. On the other hand, intestinal roll sections from Apc^Min/+^ and Apc^Min/+^*Per1^-/-^*mice exhibited aberrant crypt foci and polyps, confirming that Apc mutation induces these lesions. The H&E staining images were not significantly different between Apc^Min/+^ and Apc^Min/+^*Per1^-/-^* mice. Notably, Apc^+/-^*Bmal1^-/-^*mice reportedly display marked progression from aberrant crypt foci to locally invasive adenocarcinoma [24], although such progression was not confirmed here. Because Swiss-roll sections were prepared from 150-day-old mice, differences in disease stage among individuals at the same chronological age may have limited our ability to detect more advanced pathological progression.

In sections from *Per1^-/-^*, Apc^Min/+^, and Apc^Min/+^*Per1^-/-^* mice, β-catenin expression was observed in the intestinal villi, consistent with β-catenin is localized in normal mucosa [12]. Increased β-catenin in *Per1* deficient conditions is in line with previous findings [26] and a report using shRNA-mediated knockdown of *Per1* [27]. Furthermore, in sections of Apc^Min/+^*Per1^-/-^* mice, we also found β-catenin expression in the vicinity of the crypts, suggesting that it is possible that the loss of *Per1* induced the overexpression of β-catenin in the crypts. Western blotting further confirmed elevated intestinal β-catenin in Apc^Min/+^ mice, consistent with prior observations [11], and collectively supports a role for *Per1* in regulating β-catenin abundance. Importantly, although *Per1* deficiency increased β-catenin protein levels, Ctnnb1 mRNA levels showed no significant changes in most regions, implying that *Per1* loss primarily affects β-catenin via non-transcriptional mechanisms. Given that the Wnt/β-catenin pathway is regulated extensively at post-transcriptional and post-translational levels [28], future studies should examine β-catenin phosphorylation, subcellular localization, and components involved in β-catenin degradation.

In this study, *Per1* deficiency increased polyp number; however, the underlying mechanism cannot be concluded from our data alone. BrdU-based crypt proliferation analysis showed the Apc mutation had a significant effect, whereas *Per1* deficiency did not exert a significant effect. Thus, at least based on our assessment, the increase in polyp number caused by *Per1* deficiency on the Apc^Min/+^ background may occur without an accompanying increase in the proportion of proliferating cells within crypts. Because proliferative phenotypes in Apc^Min/+^ mucosa vary by intestinal region, age, metric, and proximity to tumors [29, 30], our Swiss-roll–based, animal-averaged approach may have masked region-specific or focal changes. Nevertheless, the observed increase in the proportion of proliferating cells attributable to the Apc mutation is consistent with previous findings. Future studies should define where *Per1* deficiency acts by quantifying Wnt target-gene mRNAs, comparing proliferative properties using organoids derived from Apc^Min/+^ and Apc^Min/+^ *Per1^-/-^* mice, and applying unbiased transcriptomic approaches such as RNA-seq.

Many studies have been conducted on the clock gene *Per1* and cancer; marked growth inhibition and apoptosis were observed when *Per1* was overexpressed in prostate cancer cells [17]. It was also found that down-regulation of the clock gene *Per1* in tumor cells accelerated cancer cell growth in vitro and tumor growth in *vivo* [31]. These previous studies suggest that *Per1* plays a role in suppressing cancer development and progression, and *Per1^-/-^* mice are more likely to develop cancer. As for the results of the present study, we found an increase in polyp number and β-catenin level due to the loss of *Per1*, which was consistent with previous studies.

## Conclusion

The present results suggest that a deficiency of the clock gene *Per1* accelerates the development and progression of colorectal cancer and that the clock gene *Per1* may play a role in inhibiting the development and progression of colorectal cancer.

## Acknowledgments

This work was supported by the Japan Society for the Promotion of Science KAKENHI (19K06360 and 24K10029 to T.J.N.) and Japan Agency for Medical Research and Development (grant number AMED-CREST 24022159 to T.J.N.). Use of AI-assisted technologies: ChatGPT (OpenAI) was used for language editing during manuscript preparation.

## Data Availability

All data for data analysis are available at Mendeley data (doi: 10.17632/tcpj358sk2.2).

## CRediT author contributions

T.N., Y.T., and T.J.N. conceived and designed the research; Y.T. T.N., and M.N. performed the experiments; T.N. and Y.T. analyzed and interpreted the data; T.J.N. contributed reagents, materials, analysis tools or data; Y.T., T.N., and T.J.N. wrote the paper.

